# Dysregulated Transferrin Receptor Disrupts T Cell Iron Homeostasis to Drive Inflammation in Systemic Lupus Erythematosus

**DOI:** 10.1101/2021.11.25.470053

**Authors:** Kelsey Voss, Arissa C. Young, Katherine N. Gibson-Corley, Allison E. Sewell, Evan S. Krystofiak, Jacob H. Bashum, William N. Beavers, Ayaka Sugiura, Eric P. Skaar, Michelle J. Ormseth, Amy S. Major, Jeffrey C. Rathmell

## Abstract

T cells in systemic lupus erythematosus (SLE) exhibit mitochondrial abnormalities including elevated oxidative stress. Because excess iron can promote these phenotypes, we tested iron regulation of SLE T cells. A CRISPR screen identified Transferrin Receptor (CD71) as important for Th1 cells but detrimental for induced regulatory T cells (iTreg). Activated T cells induce CD71 to increase iron uptake, but this was exaggerated in T cells from SLE-prone mice which accumulated iron. Treatment of T cells from SLE-prone mice with CD71 blocking antibody reduced intracellular iron and mTORC1 signaling and restored mitochondrial physiology. While Th1 cells were inhibited, CD71 blockade enhanced iTreg. *In vivo* this treatment reduced pathology and increased IL-10 in SLE-prone mice. Importantly, disease severity correlated with CD71 expression on SLE patient T cells and blocking CD71 enhanced IL-10 secretion. Excess T cell iron uptake thus contributes to T cell dysfunction and can be targeted to correct SLE-associated pathology.

## Introduction

Systemic lupus erythematosus (SLE) is a heterogenous autoimmune disease that impacts millions of people worldwide. While treatment options focus on broadly immunosuppressive drugs or B cell-targeting antibodies, SLE patients exhibit an assortment of T cell dysfunctions. These include multiple mitochondrial defects, which are hyperpolarized with elevated reactive oxygen species (ROS) and low ATP^1, 2^. Additionally, T cells in SLE can have high metabolic activity^3, 4^, an expansion of pathogenic Th1-like Th17 (Th1*) cells^5, 6^, and reduced induced regulatory T cell (iTreg) suppressive capacity. It is now evident that T cell subsets, including Th1, Th17 and Treg cells, each have unique metabolic phenotypes and requirements that offer metabolic targets to manipulate these T cell populations^7, 8, 9^. Given these distinct dependencies, metabolic targeting is a strategy of interest to treat T cells in the setting of SLE^10, 11, 12, 13^, and simultaneous inhibition of glycolysis and mitochondrial function reduced T cell-mediated inflammation in two mouse models of lupus^14^. In addition to these broad-based approaches, targeting micronutrients, such as iron, may directly impact both cellular metabolism and mitochondrial ROS. These pathways, however, remain poorly understood in SLE.

Iron is required for many cellular functions, but excess unbound labile iron can induce cellular damage via oxidative stress, mitochondrial damage, and ferroptosis. The Transferrin Receptor (CD71) is the primary iron receptor for immune cells and binds to Transferrin (Tf)-bound iron to facilitate internalization of the CD71-transferrin-iron complex. Iron flux and CD71 expression have been shown to be important for T cell activation because missense mutations in the gene for CD71, *TFRC*, result in combined immunodeficiency (CID)^15^ with defective T cell proliferation^16^. Low serum iron conditions can also impede the primary CD8 T cell response to vaccinations^17^ and Th1 responses^18^. Low iron decreased CD8 mitochondrial polarization and mTORC1 signaling, highlighting the link between iron flux, mitochondrial function, and metabolic programming in T cells. Regulation of iron metabolism may also differ among T cell subsets^19^ as Th1 cells were reported to be highly sensitive to iron depletion^20^, and CD71 blockade prevented Th17 differentiation^21^.

Although many patients with SLE have low serum iron^22^ and are often considered anemic^23^, CD4 T cells can contain higher levels of intracellular iron than controls^24^. Consistent with a role for iron in SLE, administration of hepcidin in SLE-prone mice^25^ reduced renal iron accumulation and diminished pro-inflammatory cytokine production from macrophages in the kidney^26^. Mechanisms by which iron may be dysregulated and potentially contribute to disease, however, have been unknown. Here we examined iron regulation in SLE-prone mice and patient samples. We show that SLE-prone T cells have increased CD71 expression and iron uptake that impairs normal mitochondrial physiology and mTORC1 signaling. Targeting CD71 with an antibody that promotes receptor internalization to block binding of Transferrin-iron complexes for iron uptake normalized mTORC1 signaling, mitochondrial function and metabolism. Blocking CD71 also reduced disease severity *in vivo* in SLE-prone mice and CD71 levels on SLE patient Th17 cells correlated with disease severity. These findings support a new strategy to normalize T cell metabolism and function while reducing autoimmune pathology in SLE by suppressing T cell iron uptake.

## Results

### CD71 is conditionally essential for effector T cells but suppresses iTreg

While iron metabolism may offer novel therapeutic targets with the potential to regulate T cell differentiation and mitigate mitochondrial oxidative stress, unbiased approaches have not been employed to establish the essential iron regulatory genes in T cells. We therefore designed a custom pooled library of CRISPR-Cas9 guide RNAs (gRNAs) targeting genes in iron metabolism to determine genes that are important for T cell differentiation. Naïve T cells from Cas9 transgenic mice were retrovirally transduced to express the pool of gRNAs and cells were differentiated into Th1 and iTreg cells in physiologically relevant human plasma-like media (HPLM). gRNA frequencies were determined by sequencing and comparison of guide frequencies in the resultant populations were compared to the starting pool to establish if a given gene deletion reduced or increased T cell fitness and accumulation. While CRISPR disruption of most genes had modest or no effect, deletion of *Slc25a37* (Mitoferrin-1) and *Tfrc* (Transferrin Receptor, CD71) were detrimental and led to fitness disadvantages for Th1 cells (Fig 1a). Conversely, deletion of *Tfrc* provided a strong fitness advantage to iTreg cells. *Tfrc* thus conditionally impacts T cell subsets and is essential for inflammatory T cells and inhibitory for iTreg.

**Fig. 1:**
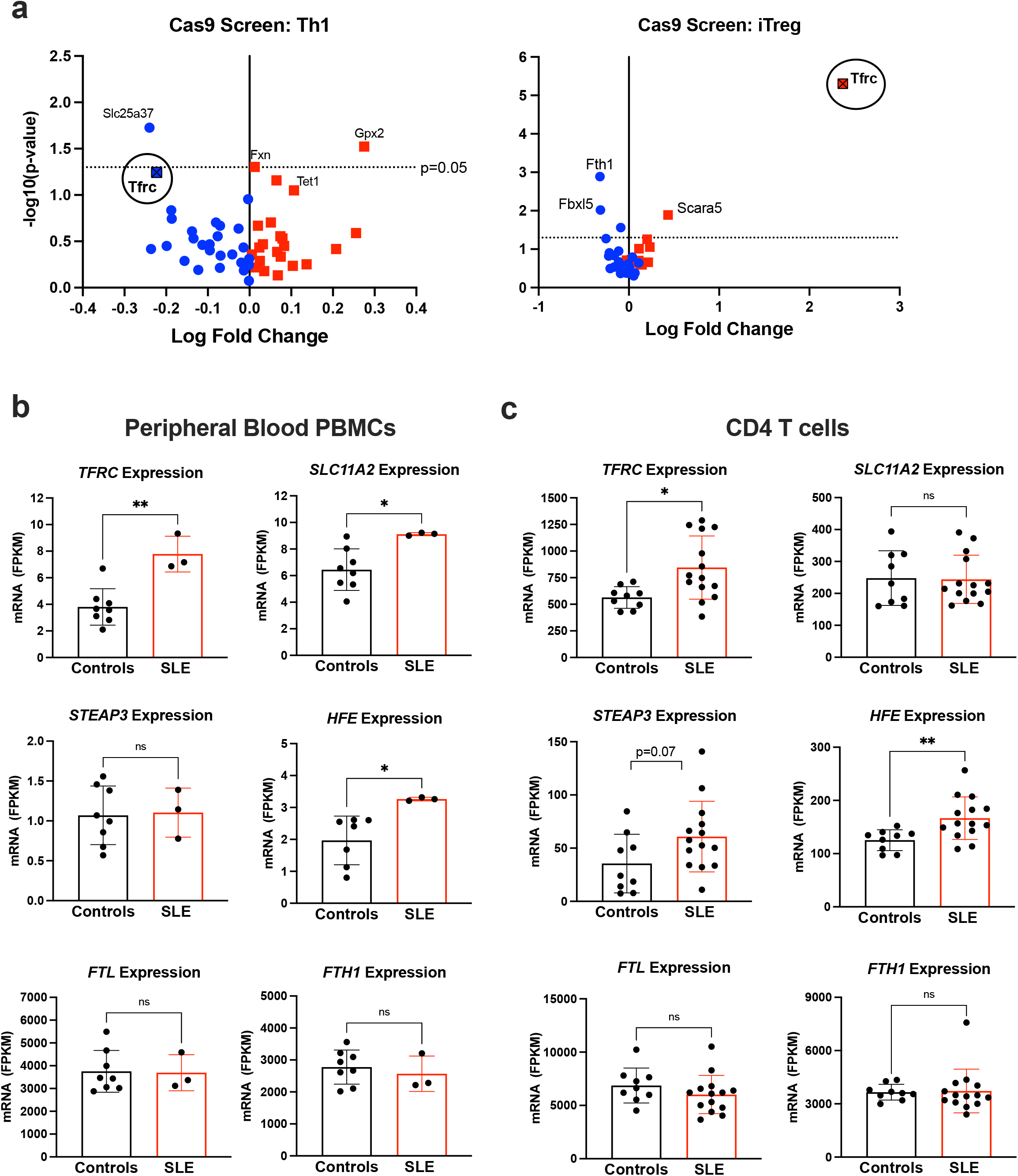
Transferrin receptor is conditionally essential for effector T cells and iTreg. **a,** Naïve CD4 T cells from Cas9-expression mice were stimulated and skewed into Th1 (left) or iTreg (right) cells. Cells were transduced with a retroviral library of guideRNAs (gRNA) specific for genes in iron metabolism or non-targeting controls. MaGeck analysis was used to determine gRNA depletion (blue) and enrichment (red) in T cell cultures over 7 days. Results are representative of two independent screens. **b, c,** Two published RNA-sequencing datasets were examined for gene expression in a subset of iron metabolism genes from patient PBMCs (b) or CD4 T cells (c). FPKM= fragment per kilobase pair per million. *P* values were determined by student’s *t*-test, two-tailed.

### Iron metabolism is dysregulated in T cells from patients with SLE

Iron metabolism in immune cells is subject to tight regulation through both systemic and cellular mechanisms to support multiple cellular functions and oxidative stress^27^. Given the increased mitochondrial abnormalities, oxidative stress, cellular metabolism of SLE T cells, we asked whether dysregulated iron handling was apparent in SLE patients in published RNA-sequencing^28^ and human genome array datasets^29^. In both SLE peripheral blood mononuclear cells (PBMCs) and purified CD4 T cells, gene expression of *TFRC* was significantly higher in SLE than healthy subjects (Fig. 1b, c). *SLC11A2*, which encodes for the divalent metal transporter 1 (DMT-1), was increased in PBMCs, but not in CD4 T cells. Six-Transmembrane Epithelial Antigen of Prostate 3 (*STEAP3*) expression, an endosomal ferrireductase, was modestly increased in CD4 T cells from SLE patients but not PBMC samples. Interestingly, hemochromatosis (*HFE*)^30^, which encodes a direct negative regulator of CD71 was significantly increased in both PBMC and CD4 T cell samples from SLE patients. Collectively, these studies suggest that CD71 and iron uptake may be dysregulated and contribute to the elevated iron concentrations and dysfunction previously measured in T cells from patients with SLE^24^.

### T cells from lupus-prone mice have altered activation and kinetics of CD71 induction

T cell receptor (TCR) activation is often impaired in T cells from patients with SLE^31, 32^. To examine the regulation of iron uptake and metabolism in lupus we examined the triple congenic mouse model of lupus-prone mice (referred to as SLE1.2.3), which closely mimics human disease^33, 34^. Naïve CD4 T cells were isolated from SLE1.2.3 mice or healthy age-matched control mice and stimulated to measure activation over time. T cells from lupus mice induced significantly lower levels of activation markers CD25 and CD69, and CD62L downregulation was delayed (Fig. 2a). In contrast, CD44 and CD71 were significantly higher in activated SLE1.2.3 T cells compared to controls (Fig. 2a, b). Additionally, cell surface CD71 remained elevated long after initial stimulation, suggesting delayed internalization or prolonged expression relative to controls that was associated with increased intracellular iron concentrations in SLE1.2.3 T cells (Fig. 2c). Therefore, both the elevated intracellular iron and altered T cell activation phenotypes were consistent between the SLE1.2.3 model and phenotypes previously demonstrated by SLE patient T cells^24^.

**Fig. 2:**
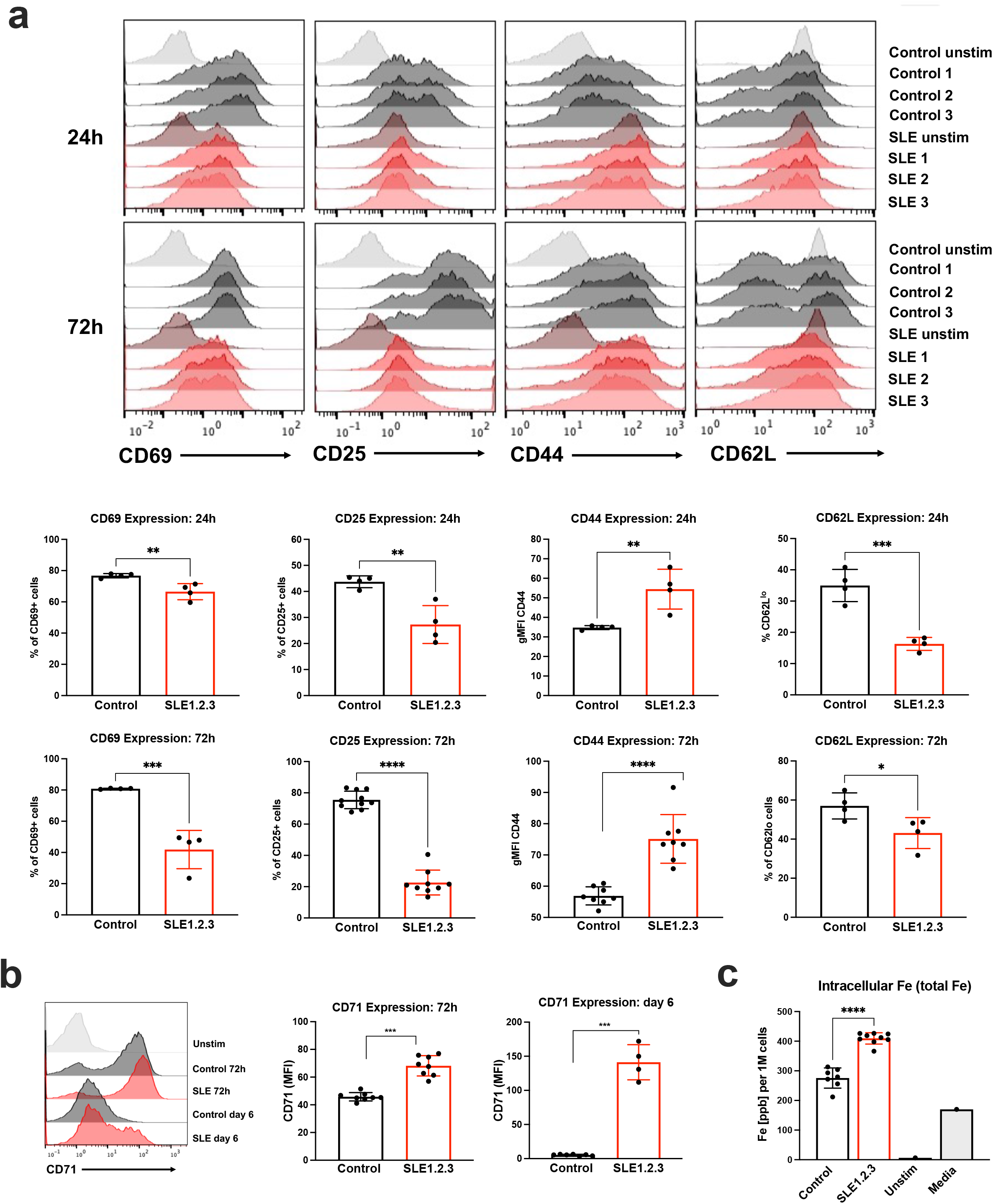
SLE1.2.3 T cells have altered T cell activation and iron regulation. **a,** Naïve T cells were isolated from SLE1.2.3 mice or age matched healthy control and stimulated with splenocytes + anti-CD3. Activation markers CD69, CD25, CD44, and CD62L were measured by flow cytometry at 24h and 72h post-activation. **b,** CD71 expression at 72h and 6 days post-activation with a representative plot (left). **c,** T cell pellets from 72h post-activation were subjected to ICP-MS to determine intracellular iron concentrations. ‘Unstim’ are T cells that were left unstimulated and ‘Media’ is a control for iron content in the cell culture media only. Ppb= parts per billion. (a,b,c) Significance was determined by student’s two-tailed *t*-test, representative of 3 independent experiments.

### CD71 blockade reduces iron flux and normalizes SLE T cell activation in vitro

Anti-CD71 antibodies have been reported to bind CD71 and block association with Transferrin-iron complexes and stimulate receptor internalization without iron uptake^35, 36^. Consistent with receptor blockade and/or internalization, a large decrease in cell surface CD71 expression was observed by flow cytometry in activated T cells when cells were treated with CD71 blocking antibody compared to the isotype control (Fig. 3a). Decreased available CD71 correlated with significantly decreased intracellular labile iron (Fig. 3b). Because CD71 expression and iron flux are important for T cell activation, we tested whether anti-CD71 would alter T cell activation. Surprisingly, CD69 upregulation was restored to SLE1.2.3 T cells treated with anti-CD71 whereas control T cells were unaffected (Fig. 3c). Conversely, CD44 overexpression in SLE1.2.3 T cells was lowered to normal by anti-CD71 treatments (Fig. 3d). Furthermore, T cells in patients with SLE have a well-documented deficiency in IL-2 production^37^, and the percentage of IL-2-producing T cells was increased in SLE1.2.3 T cells activated with anti-CD71 (Fig. 3e).

**Fig. 3:**
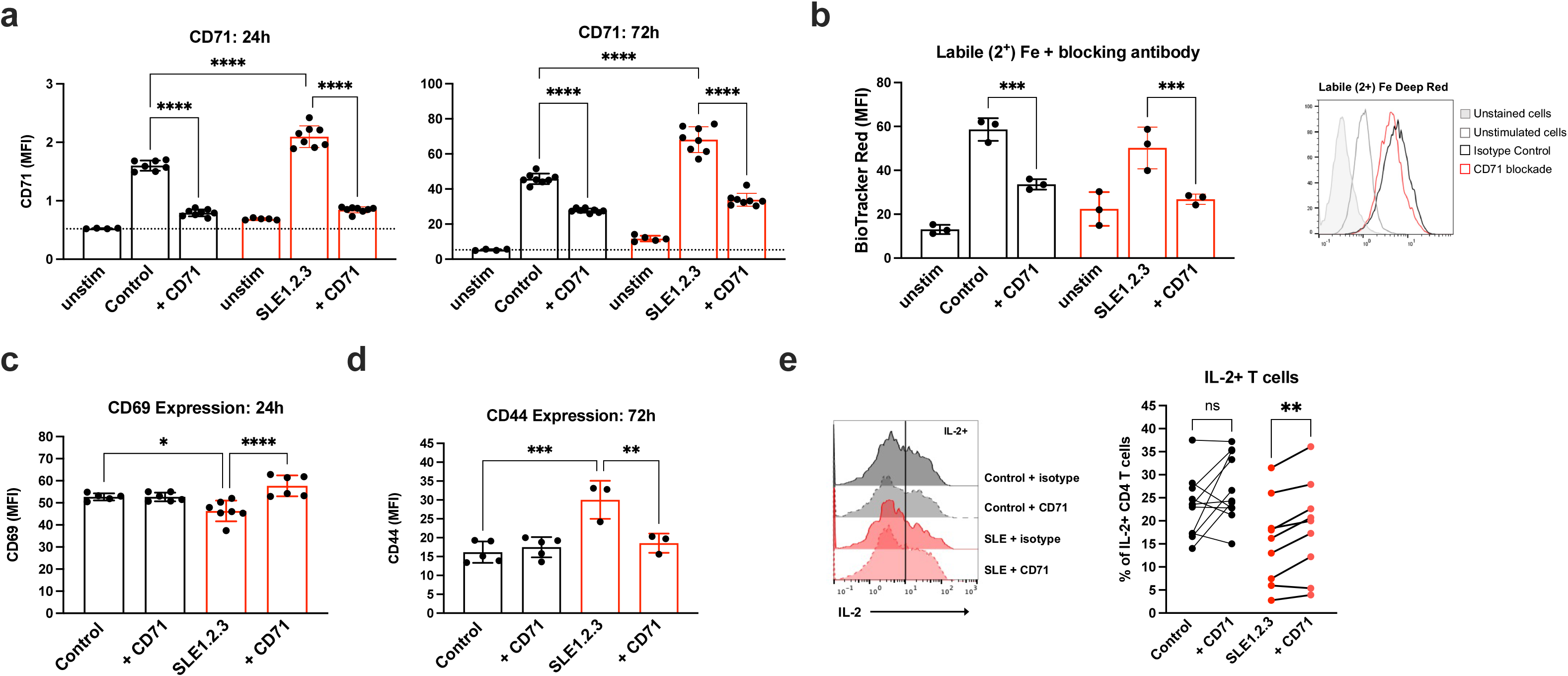
CD71 blockade normalizes T cell activation and IL-2 production in SLE1.2.3 T cells. **a,** Naïve T cells were activated in the presence of an isotype control antibody or CD71 blocking antibody. CD71 expression was measured by flow cytometry at 24h (left) and 72h (right) post-activation. **b,** Labile iron in T cell cultures activated as in (a) was measured with BioTracker Red dye at 72h post-activation, representative plot shown (right). Similar results were obtained in 3 independent experiments. **c,** CD69 expression was determined by flow cytometry 24h post-activation in T cell cultures as in (a). **d,** CD44 expression was determined by flow cytometry 72h post-activation. **e,** T cell cultures were restimulated on day 5 post-activation with PMA/ionomycin to quantify the percentage of IL-2+ CD4 T cells. (a-d) One-way ANOVA with Sidak’s multiple comparisons test. (e) Paired ANOVA with Sidak’s multiple comparisons test, results are from 3 experiments combined.

### CD71 blockade restores mTORC1 signaling and mitochondrial function in SLE-prone T cells

To determine the mechanism by which targeting CD71 targeting may rescue functional abnormalities in SLE T cells, we first tested if this approach affected the mTORC1 pathway. mTORC1 integrates the status of multiple cell nutrient and metabolic conditions with cell signaling to phosphorylate targets such as p70S6 Kinase and promote anabolic metabolism for effector T cells. High levels of mTORC1, however, can be detrimental to Treg. Indeed, a reduction in iron flux by anti-CD71 significantly reduced p-S6 in lupus T cells (Fig. 4a), consistent findings that iron chelation suppresses mTORC1 activation^38, 39^. Because mTORC1 broadly regulates anabolic metabolism the effects of CD71 blockade were tested on T cell size, which was significantly reduced and normalized in SLE1.2.3 T cells with anti-CD71 (Fig. 4b).

**Fig. 4:**
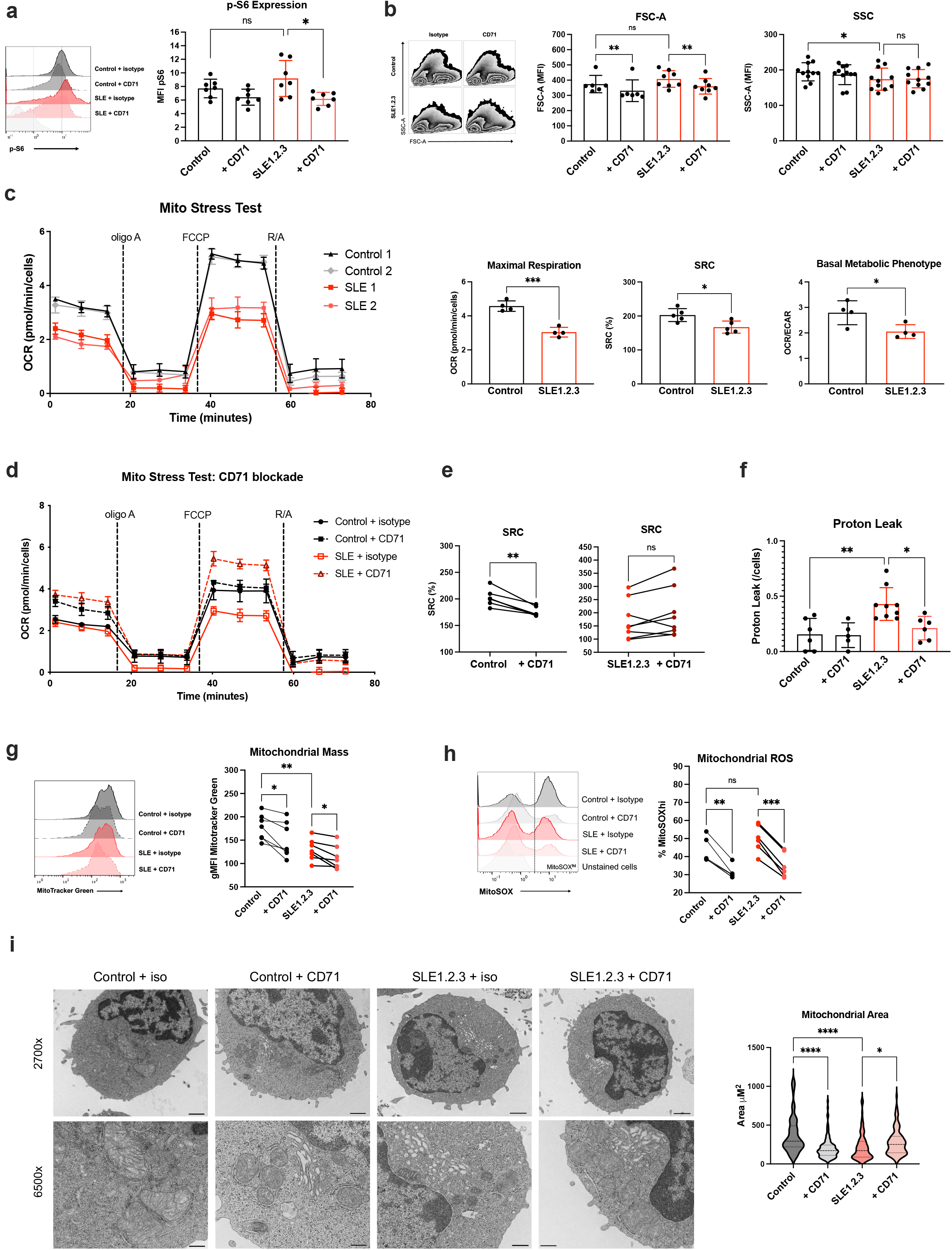
CD71 blockade restores metabolic and mitochondrial function in SLE1.2.3 T cells. **a,** T cell cultures on day 5 of stimulation were analyzed by intracellular flow cytometry for phospho-S6. Results from two independent experiments. **b,** Forward scatter and side scatter (FSC, SSC) were measured in T cell cultures. Results from three independent experiments. **c,** Extracellular flux analysis on day 5 post-activation. Oxygen consumption rate (OCR) during a Mito Stress Test is shown from two biological replicates (left). Maximal respiration and spare respiratory capacity (SRC) were quantified from Mito Stress Test. Basal OCR from each sample as in (a) was divided by the basal extracellular acidification rate (ECAR) to determine the relative energic metabolic phenotype of each sample (right). R/A= Rotenone and antimycin A. **d,** Naïve T cells were activated in the presence of CD71 blocking antibody or an isotype control and subjected to Mito Stress Test on day 5 post-activation. Representative plot of 4 independent experiments. **e,** SRC (%) calculated from (b). **f,** Proton leak calculated from (b). **g,h,** Mitotracker green (e) and MitoSOX Red (f) staining was measured by flow cytometry on day 5 post-activation. **i,** T cell cultures were processed for EM. A minimum of 100 mitochondria were analyzed in each sample group to quantify mitochondrial area (right). Scale bar at 2700x= 1μM, scale bar at 6500x= 400nm. (a,b,f,g,h,i) One-way ANOVA with Sidak’s multiple comparisons test (c) Two-tailed student’s *t*-test (e) Paired *t*-test.

To test if CD71 blockade also corrected mitochondrial defects in SLE1.2.3 T cells, a baseline of mitochondrial function in SLE1.2.3 T cells was established by extracellular flux analyses. T cells from SLE1.2.3 mice had significantly lower mitochondrial respiration than controls, both at baseline and at maximal respiration (Fig. 4c). SLE1.2.3 mitochondria also had a lower spare respiratory capacity (SRC). While extracellular acidification rate (ECAR) was unchanged (Fig. Supp. 1a), SLE1.2.3 T cells had a basal bioenergetic profile reflective of reduced oxidative respiration relative to control T cells.

Decreased mitochondrial respiration in SLE1.2.3 T cells may have been due to reduced mitochondrial quality caused by increased iron loads. All T cells required some level of iron because T cells incubated with the cell permeable iron chelator CPX had reduced mitochondrial respiration regardless of disease status (Fig. Supp. 1b). In contrast, selective targeting of CD71-mediated iron uptake with an anti-CD71 blocking antibody increased mitochondrial respiration in SLE1.2.3 T cells (Fig 4d). Although basal metabolism was not significantly affected (Fig. Supp. 1c), SLE1.2.3 T cells showed a trend to increase spare respiratory capacity (SRC) upon CD71 blockade while this was decreased in control T cells (Fig. 4e). SLE1.2.3 T cells also had a significantly higher mitochondrial proton leak, a measure of electron transport efficiency across the mitochondrial membrane outside of ATP synthase and mitochondrial quality^40^, than control cells (Fig. 4f). Importantly, proton leak was restored in SLE1.2.3 T cells to levels of controls by anti-CD71.

Mitochondrial mass was reduced in SLE1.2.3 T cells relative to control T cells (Fig. 4g). Interestingly, anti-CD71 treatment resulted in a further decrease in mitochondrial mass as determined by Mitotracker Green staining in both control and SLE1.2.3 T cells. CD71 blockade also resulted in a large decrease in mitoROS in both control and SLE1.2.3 T cells (Fig. 4h). The ultrastructure of the mitochondria was next analyzed by electron microscopy (EM) and consistent with less mitochondrial mass when measured through dye accumulation, SLE1.2.3 T cell mitochondrial cross-sectional area was significantly lower than controls (Fig. 4i). Intriguingly, while anti-CD71 treatment reduced mitochondrial area in control T cells, anti-CD71 treatment increased and normalized mitochondrial area in SLE1.2.3 T cells.

### CD71 differentially impacts T cell subsets

CRISPR/Cas9 screening indicated conditional essentiality of *Tfrc* in Th1 but not iTreg cells (Fig 1a). Therefore, we tested whether anti-CD71 exerted distinct impacts on different CD4 T cell subsets. Naïve T cells from control mice were activated with CD71 blockade or an isotype control during differentiation for Th1, Th17, and iTreg cells. Anti-CD71 treatment led to increased apoptotic cell death in Th1 cultures but had limited or no effect on iTreg or Th17 cultures (Fig. 5a). Mechanistically, this increased apoptosis was associated with extensive DNA damage as indicated by phosphorylation of H2AX, a hallmark of double-stranded breaks (Fig 5b).

**Fig. 5:**
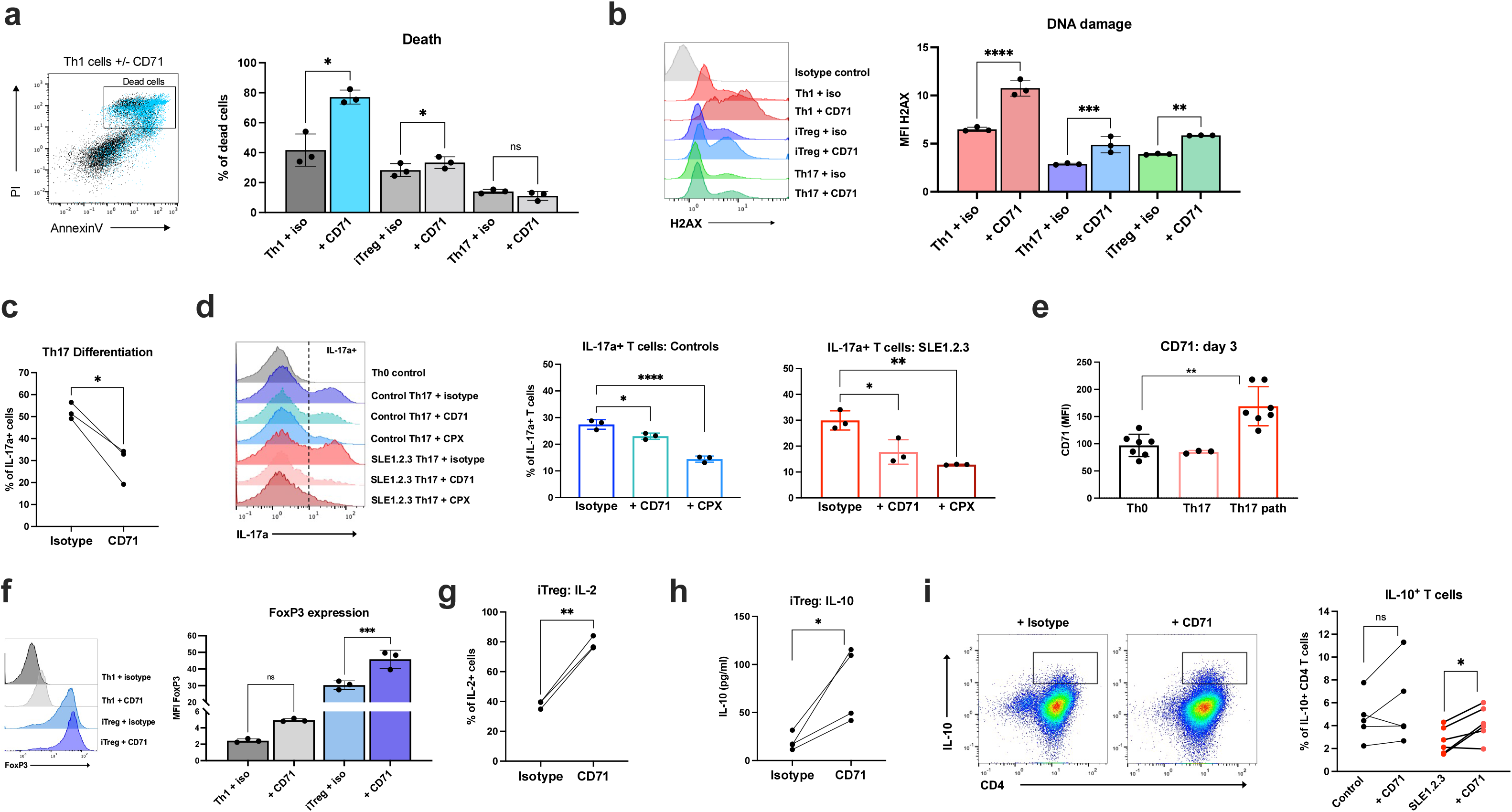
CD71 blockade differentially impacts T cell subsets. **a,** Cell death was measured by AnnexinV and propidium iodide (PI) staining in flow cytometry. A representative plot of Th1 cells is shown (left). **b,** T cell cultures as in (a) were analyzed for phosphorylated H2AX as an indicator of DNA damage. **c,** Th17 cultures were restimulated with PMA/ionomycin on day 5 to determine the percentage of IL-17a+ cells with flow cytometry. **d,** Naïve CD4 T cells from SLE1.2.3 mice or B6 controls treated with either anti-CD71 or CPX. IL-17+ cells determined as in (c). **e,** CD71 expression on activated T cells with no cytokines (Th0), Th17 cytokines (TGFβ + IL-6) or pathogenic Th17 cytokines “Th17 path” (TGF-β + IL-6 + IL-23 + IL-1β). Results are representative of three independent experiments. **f,** iTreg and Th1 cultures +/- CD71 blockade were analyzed for FoxP3 expression on day 5 post-activation. **g,** iTreg cultures were analyzed for IL-2 production as in (c). **h,** Naïve T cells were subjected to iTreg differentiation for 3 days. Supernatants were analyzed for IL-10 by ELISA and normalized to cell number. **i,** Naive CD4 T cells isolated from SLE1.2.3 and control mice. Day 4 cells were restimulated as in (c) to determine percentage of IL-10+ T cells.. (a,b,d,f,i) One-way ANOVA with Sidak’s multiple comparisons test. (c,g,h) Paired student’s *t*-test.

Despite a lack of effect on the viability of Th17 cells, anti-CD71 treatment interfered with the induction of this inflammatory T cell subset. The percentage of IL-17+ cells was reduced with CD71 blockade compared to controls (Fig 5c). Furthermore, CD71 blockade during early activation and differentiation interfered with Th17 differentiation (Fig. 5d). Th17 cells treated with CPX had a further reduction in IL-17 production in both control and SLE1.2.3 T cells, suggesting that both acute iron availability and iron loads during early T cell differentiation regulate IL-17 production and Th17 identity. Patients with SLE often exhibit an expansion of pathogenic Th1-like Th17 (Th1*) cells that secrete both IFN-γ and IL-17^6^. Interestingly, Th17 cultures skewed in pathogenic Th1/Th17 conditions expressed higher levels of CD71 after activation compared to both Th0 (non-differentiated) and non-pathogenic Th17 cells (Fig. 5e). Together, these data support a role for increased CD71 and iron flux for both Th17 differentiation and the pathogenicity of Th17 cells.

The effects of anti-CD71 were next examined specifically on iTreg cells. Consistent with enhanced iTreg fitness and accumulation after CRISPR deletion of *Tfrc* (Fig. 1a), anti-CD71 significantly increased FoxP3 expression in iTreg cells compared to the isotype control (Fig. 5f). The percentage of iTreg cells producing IL-2 also doubled, and the amount of IL-10 secreted into cell supernatants in iTreg conditions was increased with anti-CD71 treatment (Fig. 5g, h). Finally, naïve T cells isolated and differentiated into iTreg cells produced significantly more IL-10 with anti-CD71 treatment (Fig. 5i). These data show that CD71 blockade can modulate T cell differentiation to suppress effector T cells and boost FoxP3 expression and IL-10 production of iTreg cells.

### CD71 blockade reduces pathology and autoimmunity in SLE1.2.3 mice

Based on the ability of CD71 blockade to improve the activation and mitochondrial functions of SLE1.2.3 T cells, we tested if this treatment could reduce *in vivo* markers of autoimmunity in lupus mice. To this end, 4 to 4.5-month-old SLE1.2.3 mice and age-matched B6 controls were treated with isotype control or anti-CD71 antibody twice a week for 4 weeks (Fig. 6a). SLE1.2.3 mice with pre-established autoimmunity determined by anti-dsDNA IgG antibodies at the start of the treatment course were included in the study (Fig. 6b). By the end of the treatment course, anti-dsDNA antibodies in SLE1.2.3 mice treated with anti-CD71 were reduced compared to the isotype control group (Fig. 6c). Anti-nuclear antibodies (ANA) and anti-histone antibodies also showed decreased trends (Fig. Supp. 2a, b). Spleens and lymph nodes (LNs) were harvested and CD4 T cells isolated, revealing a significant drop in CD4 T cell numbers in anti-CD71-treated mice (Fig. 6d). Of note, splenomegaly showed a trend to reduce (Fig. Supp. 2c), but total B cell numbers appeared unaffected (Fig. Supp. 2d). Treatment was well-tolerated, and no signs of anemia were detected by complete blood count (CBC) analysis (Fig. Supp. 2e-g). Analysis of activation markers immediately *ex vivo* showed SLE1.2.3 T cells exhibited much higher CD69 expression which was not impacted by the anti-CD71 treatments (Fig. Supp. 2h). There was, however, a rescue in the intensity of CD44 expression of the anti-CD71-treated group in SLE1.2.3 T cells (Fig. 6e).

**Fig. 6:**
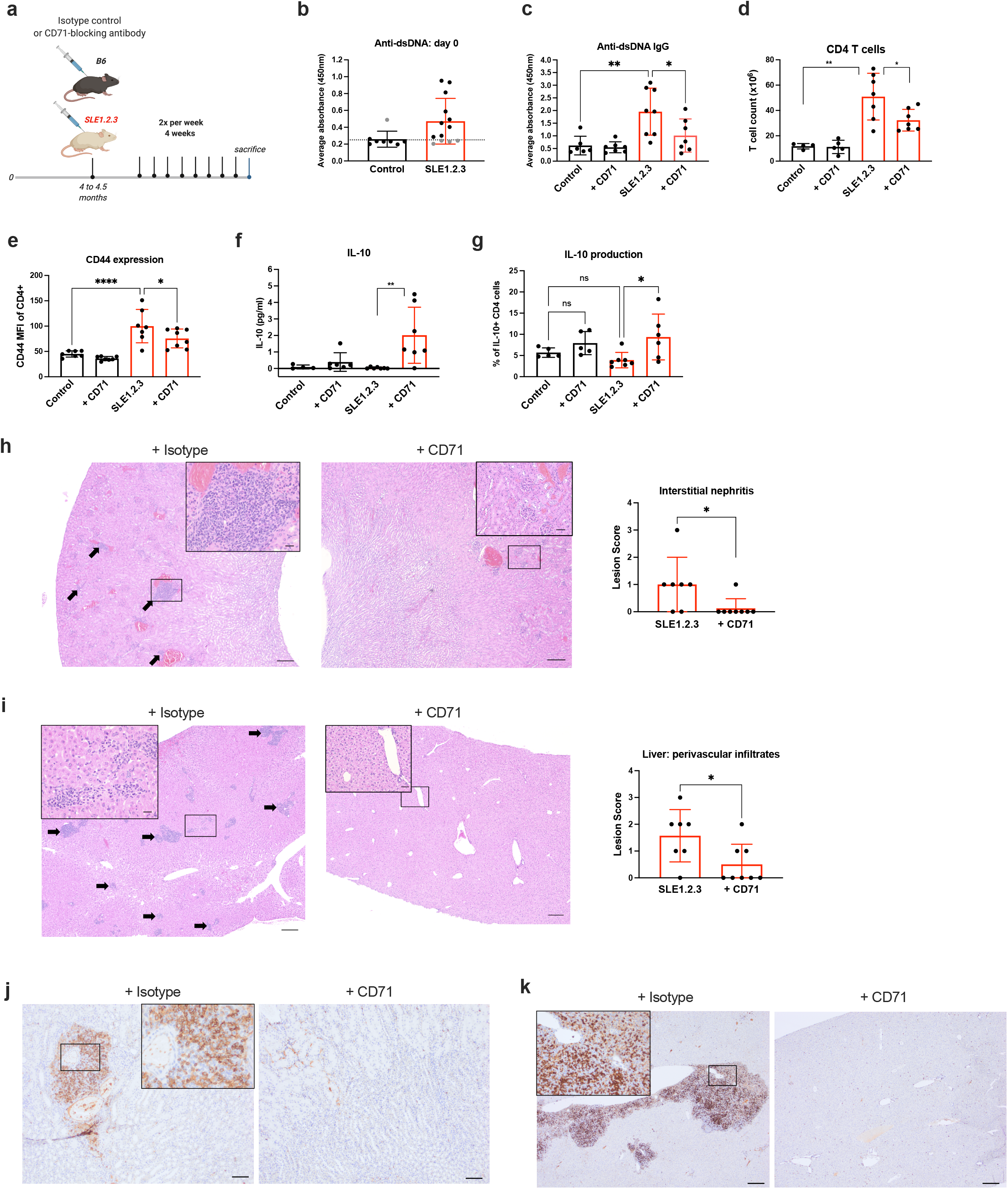
CD71 blockade reduces autoimmunity and pathology in SLE1.2.3 mice. **a,** SLE1.2.3 mice or age-matched B6 controls were treated twice per week with an isotype control or anti-CD71 for 4 weeks. **b,** Anti-dsDNA antibodies were measured in serum before the treatment regimen. The dotted line represents the cut-off value for inclusion criteria. **c,** Endpoint serum levels of anti-dsDNA Ig antibodies. **d,** CD4 T cells were isolated from spleen and lymph nodes by negative selection and cell counts determined by automated cell counter. **e,** CD4 T cells as described in (d) were stained for CD44 and examined by flow cytometry. **f,** Endpoint sera were measured for IL-10 concentrations. **g,** CD4 T cells from spleens and lymph nodes were stimulated with PMA/ionomycin. IL-10+ CD4 T cells were quantified by flow cytometry. **h,i,** Pathological assessment of inflammation in the kidney (h) and liver (i) was scored semi-quantitatively from hematoxylin and eosin (HE) stained tissue sections. Arrows indicate infiltrates of lymphocytes and plasma cells. Large scale bar= 200μm, inset bar=20μm. **j,k,** Representative IHC staining of CD4 T cells in the kidney (j) and liver (k) is shown. All experiments were N=3. (c,d,e,f,g) One-way ANOVA with Sidak’s multiple comparisons test. (h,i) Two-tailed student’s *t*-test.

Consistent with increased IL-10 secretion from iTreg cultures treated with anti-CD71 (Fig. 5h) there was a large increase of IL-10 in the serum of SLE1.2.3 mice with anti-CD71 treatments (Fig. 6f). Furthermore, the percentage of CD4 T cells producing IL-10 was significantly increased (Fig. 6g), suggesting that CD71 targeting may rewire CD4 T cells *in vivo* to secrete IL-10. We hypothesized that Treg may be a source of the IL-10 as our CRISPR screen and CD71 blockade in vitro was beneficial to iTreg cultures (Fig. 1a). Although absolute numbers of natural Tregs (nTreg) did not increase significantly, nTreg frequencies trended upward in SLE1.2.3 mice treated with CD71 antibody (Fig. Supp. 2i). Conversely, the frequency of T follicular regulatory (Tfr) cells showed a decreased trend in anti-CD71-treated mice that may be consistent with reduced autoimmunity and potential lessened germinal center activity (Fig. Supp. 2j).

Kidneys and livers were next analyzed to assess SLE-associated pathology and determine if CD71 blockade prevented tissue damage. Histopathologic assessment of inflammation was blindly scored with a semi-quantitative inflammation scale in stained tissue sections^41^. In the kidney, interstitial nephritis and glomerular loop thickening were evident in the kidneys of SLE1.2.3 that received isotype control antibody (Fig. 6h). These lesions were significantly reduced with anti-CD71 treatments. SLE1.2.3 mice also exhibited periportal infiltrates of lymphocytes and plasma cells in the liver (Fig. 6i) that were greatly reduced with anti-CD71 treatment. CD4 T cell immunoreactivity was noticeably reduced by anti-CD71 treatment in the kidney (Fig. 6j). Additionally, periportal lymphocytes stained robustly for CD4 and this was also reduced in the anti-CD71 treatment group (Fig. 6k), demonstrating that anti-CD71 treatments can significantly reduce CD4 T cell infiltration and autoimmune tissue damage. We also found macrophages were abundant in the liver and kidney lesions and this decreased with anti-CD71 treatment (Fig. Supp. 2k).

### CD71 expression on T cells corresponds to disease activity and increased Th17 cells in SLE patients

T cells from healthy donors and SLE patients were next evaluated for dependence on CD71. Naïve human T cells were activated from three healthy donors in either undifferentiated Th0 conditions or Th17 conditions. Th17-skewed T cells from each donor contained significantly higher intracellular iron concentrations than Th0 cells (Fig. 7a) consistent with a role for iron in Th17 differentiation in human T cells. Therefore, T cells from patients with SLE and healthy donors with no history of autoimmune conditions were collected to characterize the expression and role of CD71 (**Table 1 and** Fig. Supp. 3a). In both control and patient samples there was a notable correlation between the percentage of Th17 cells and the level of CD71 expression on all CD4 T cells (Fig. 7b), consistent with a relationship between CD71 expression and Th17 cell differentiation. On average, CD71 expression was increased on CD4 T cells from patients with SLE compared to controls (Fig. 7c). Patients with SLE also had higher percentages of Th17 cells in their CD4 T cell compartment and CD71 expression on these cells was elevated in many patients (Fig. 7d, e). Despite increased CD71 and CD44 expression (Fig. Supp. 3b), CD69 did not significantly differ in patients with SLE compared to controls to suggest that acute activation marker status alone cannot explain the increased CD71 expression. Furthermore, although SLE patients often develop anemia, CD71 expression on T cells did not negatively correlate with hemoglobin levels (Fig. Supp. 3c) suggesting that anemia does not dictate CD4 T cell expression of CD71. Patients with high CD71 expression may represent a subset of SLE with more active disease and Th17 cell involvement. SLE disease activity assessed by Systemic Lupus Erythematosus Disease Activity Index (SLEDAI) scores showed a trend towards patients with worse disease to have higher CD71 expression on CD4 T cells (Fig 7f), although there were no apparent differences in the frequency of Th17 cells (Fig. 7g). CD71 expression on Th17 cells, however, did significantly correlate with disease severity (Fig. 7h).

**Fig. 7:**
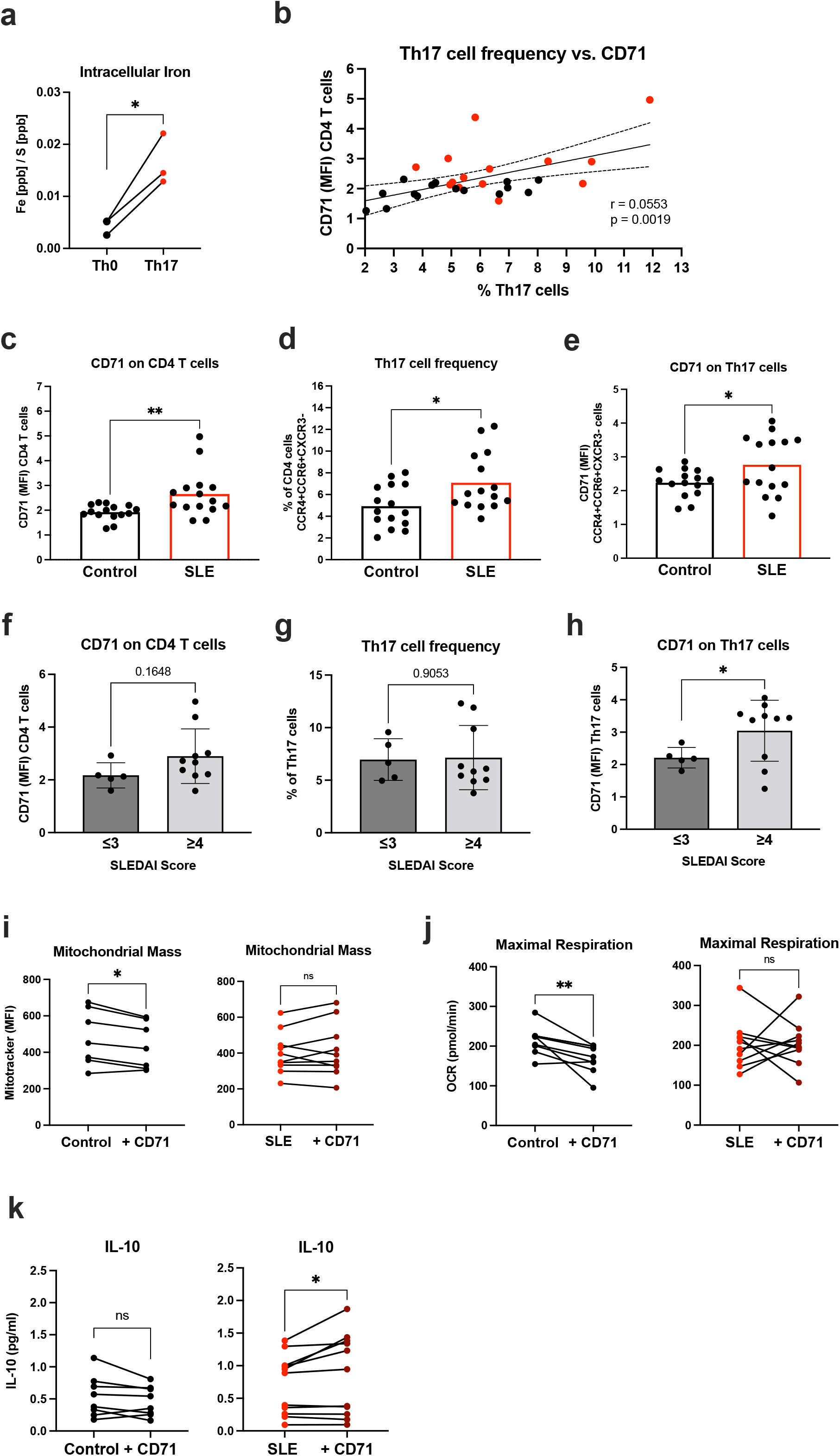
CD71 expression correlates with Th17 cells in lupus patients. **a,** Naïve human T cells were activated for 5 days in non-differentiating conditions (Th0) or Th17 conditions. Fe measured by ICP-MS and normalized to sulfur (S) for protein content. Paired Student’s t-test. **b,** PBMCs were isolated from healthy donors (Control) and SLE patients (SLE). Percentage of Th17 cells in CD4 T cell compartment compared to CD71 expression on bulk CD4 T cells. Controls=black, SLE= red. Pearson correlation was used to determine r value and statistical significance. **c,** CD71 expression on bulk CD4 T cells. **d,** Percentage of CD4 T cells defined as Th17 cells. **e,** CD71 expression on Th17 cells. **f,g,h,** Flow cytometry data from patients with SLEDAI scores of 3 or less compared to 4 or higher. **i,** Mitotracker Green was quantified by flow cytometry on day 4 post-activation. Paired t-test. **j,** Maximal respiration was determined from Mito Stress Test assay on day 4-5 post-activation. **k,** IL-10 concentrations in day 3 supernatants measured by ELISA. (c-h) Student’s two-tailed *t*-test. (i,j,k) Paired student’s *t*-test.

### Targeting CD71 in SLE patient T cells increases IL10 secretion

To test the effects of CD71 targeting in lupus patient T cell samples, we employed a human-specific CD71 blocking antibody mirroring our previous tests with a murine-specific antibody in SLE1.2.3 mice. The antibody reduced iron uptake in activated T cells (Fig. Supp. 3d). In agreement with CD71 blockade in murine T cells, anti-CD71 did not interfere with activation of human T cell samples as measured by activation markers CD25 and CD44 (Fig. Supp. 3e). Anti-CD71 led to a drop in mitochondrial mass in healthy control but not SLE patient T cell samples (Fig. 7i).

Similarly, analyses of mitochondrial respiration demonstrated a consistent drop in maximal respiration in control T cell samples (Fig. 7j), while SLE patient T cells had heterogeneous responses, and many showed increased maximal respiration. Despite this heterogeneity, CD71 blockade led to a significant increase in IL-10 secretion from SLE patient T cells (Fig. 7k). Altogether, CD71 targeting in SLE patient T cell cultures appeared to restore mitochondrial function and boosted IL-10 production in a subset of samples.

## Discussion

SLE is a complex autoimmune disease with a multitude of factors contributing to its etiology including environment, genetics, and immune cell dysregulation. Among these, iron metabolism has recently been recognized as a potential driver of disease. Dietary iron can aggravate lupus symptoms in patients with SLE^42^, and there remains an unmet need to understand how iron metabolism influences immune cell function in the setting of lupus^43^. While iron has been suggested as a target in inflammatory diseases, immune cell function studies of iron metabolism have focused mainly on innate immune cells such as macrophages. Lowering serum iron levels in mouse models of lupus has shown promising reductions in autoimmune pathology, but these efforts have not focused on T cell phenotypes and the differential effects of iron on T cell subsets^26, 44^. Furthermore, mechanisms by which iron regulates T cell function and differentiation have been unclear.

Here we show that targeting iron metabolism via CD71 can reduce disease manifestations of SLE through T cell regulation by multiple mechanisms. First, blocking CD71 reduced iron flux and normalized T cell activation. While some markers of T cell activation in SLE T cells were reduced upon TCR stimulation, CD71 induction was heightened and remained elevated long after initial stimulus. This increased capacity for iron uptake likely drove the accumulation of iron in SLE T cells despite the predisposition for SLE patients to develop low serum iron. High levels of intracellular iron, in turn, impaired mitochondrial respiration and quality control to promote ROS and reduce ATP production efficiency. Indeed, mitochondrial activity, mass, and mitoROS were also rescued in activated SLE T cells. Second, mTORC1 is a nutrient sensitive kinase complex and master regulator of anabolic growth processes and was also regulated in a CD71-dependent manner. Elevated mTORC1 activity can promote effector T cells, including Th17, and suppress Treg function and stability, and may contribute to the finding of increased Th17 and decreased Treg in SLE patients.

Our study also sheds new light on iron biology in different T cell subsets and differentiation. First, using both an unbiased genetic and antibody-blocking approach, iTreg cells thrived in CD71-targeting conditions whereas Th1 cultures had increased levels of cell death. Second, T cells produced more IL-2 and IL-10, suggesting that iron metabolism can be manipulated to control regulatory cytokine activity. In agreement with this, iron limitation in Jurkat T cells increased IL-2 secretion^45^. CD71 targeting was particularly beneficial to iTregs by increasing FoxP3 expression and suggests that they could have enhanced suppressive capacity. Indeed, Tregs may have an increased susceptibility to iron-induced oxidative stress and/or ferroptosis that depends on glutathione peroxidase 4 (Gpx4) for cellular redox homeostasis. Interestingly, a loss of Gpx4 activity in Tregs resulted in IL-1β release during ferroptosis that promoted IL-17a production in CD4 T cells^46^.

We also show that CD71 expression is associated with Th17 cell development in humans and that Th17 cells cultured in in physiological media acquire more intracellular iron than undifferentiated, activated T cells from the same donor. Because many lupus patients exhibit pathogenic Th1* cells, one strategy for treating Th17-driven inflammation is to target metabolic regulators of the conversion of non-pathogenic Th17 cells to pro-inflammatory pathogenic Th17 cells^47, 48^. Pathogenic Th17 cells expressed higher levels of CD71 than non-pathogenic Th17 cells, and this may be mediated in part by IL-1β. In support of this, IL-1 was required for the induction of pathogenic Th17 cells in a model of experimental autoimmune encephalopathy (EAE)^49^. Additionally, iron-deficient mice failed to develop EAE^50^ and a CD4 T cell-specific deletion of *Tfrc* also failed to drive Th17 neuroinflammation via decreased GM-CSF mRNA stability^51^. Regulators of iron metabolism may therefore reveal novel regulators of Th17 cell pathogenicity and together, these data demonstrate a strong association between sustained CD71 expression, iron accumulation, and development of a pathogenic Th17 program.

*In vivo*, CD71 blockade was well-tolerated, significantly reduced autoantibodies, and prevented disease-specific pathology and CD4 T cell infiltration into the kidneys and liver. A surprising finding was that blocking CD71 increased systemic IL-10 levels in the serum and IL-10-producing T cells. Although IL-10 is generally thought to be anti-inflammatory role, its role in SLE is controversial, and has been associated with more active disease^52^. This may be due to compensation for chronic inflammation as B cells in SLE mice treated with IL-10 blocking antibodies produced less anti-dsDNA antibodies. Here we found SLE1.2.3 mice had a significant reduction in anti-dsDNA titers that correlated with increased systemic IL-10 upon anti-CD71 treatment^53^. IL-10 was also linked to increased apoptosis of SLE patient CD4 T cells by inducing Fas and FasL expression^54^. However, recent work investigating the dual role of IL-10 in murine lupus showed that blocking the IL-10 receptor *in vivo* accelerated disease and immune dysregulation^55^.

Targeting CD71 may provide a new therapeutic opportunity in SLE. An antibody blockade treatment used here in SLE1.2.3 mice to induce CD71 internalization and prevent binding to Transferrin-bound iron. This was well-tolerated here and in previous studies^56^ and no unintended anemia was observed. Anti-CD71 treatments have also been investigated in humans for cancer therapy to block the CD71 iron transport function in cancer cells^36^. However, T cell cultures from SLE patients were heterogeneous and many responded differently than control T cells to CD71 blockade during activation. Despite a small sample size, there appeared to be a subset of SLE patients with high CD71 expression on Th17 cells which correlated with a worse SLEDAI score. It is likely that anti-CD71-induced changes on this population may be concealed in the bulk cultures analyzed here. Additionally, given the benefits to iTreg populations, CD71 targeting could also benefit patients that have a low Th17/Treg ratio by boosting nTreg function and simultaneously depleting Th1* cells. These data suggest CD71 is a rational target to further explore for SLE T cells. Interestingly, other autoinflammatory Th17 diseases display iron deposits in affected tissues such as the central nervous system of multiple sclerosis (MS) patients^57^ and the synovial fluid of rheumatoid arthritis (RA) patients^58^, suggesting these findings may be relevant to inflammatory diseases beyond lupus.

## Methods

### Mice

All experiments were performed at Vanderbilt University with Institutional Animal Care and Utilization Committee (IACUC)-approved protocols. Cages were maintained in a pathogen-free facility with ventilated cages and *ad libitum* food and water. For anti-CD71 experiments, control C57BL/6 (Jax #000664) or age-matched lupus-prone SLE1.2.3 (Jax # 007228) mice were treated with 200μg of CD71 blocking antibody (clone R17 217.1.3/TIB-219) or an isotype control antibody starting at 4-4.5 months of age (BioxCell). Treatment was administered twice a week via IP injection for 4 weeks.

At the time of euthanasia, a terminal blood draw was performed, and complete blood count (CBC) analysis was obtained using the Forcyte (Oxford Science) hematology analyzer. A full gross examination and tissue collection was performed by a board-certified veterinary pathologist and tissues were immersion fixed in 10% neutral buffered formalin for approximately 3 days before routine processing and embedding in paraffin. Sections were cut at 5µm and stained with HE for analysis.

### Patient samples

Blood samples were obtained from patients with SLE and control subjects enrolled in the Inflammation, Cardiovascular Disease, and Autoimmunity in Rheumatic Diseases (ICARD) Study at Vanderbilt University Medical Center. For inclusion in the study all subjects were required to be 18-years of age or older. Patients required a rheumatologist diagnosis of SLE for inclusion and all patients with SLE met the 1997 American College of Rheumatology revised classification criteria for SLE^59^. Control subjects were excluded if they had an inflammatory autoimmune disease. Vanderbilt University Medical Center Institutional Review Board approved the study (IRB# 150544) and all subjects provided written informed consent. Clinical information was gathered via chart review by a rheumatologist. Peripheral blood was collected in heparin-coated tubes. PBMCs were then isolated by Ficoll gradient separation, subjected to flow cytometry, and stored in liquid nitrogen.

Two panels of antibodies were run on patient PBMC samples. All antibodies were purchased from BioLegend. The first panel was used to quantify CD71 and activation markers on bulk CD4 T cells. CD3-APC/Cy7 (clone HIT3a), CD4-PE/Cy7 (clone a161A1), CD25-BV605 (clone BC96), CD44-eFluor450 (clone IM7), CD69-FITC (clone FN50), and CD71-APC (clone CY1G4). The second panel was used to quantify Th17 cells and CD71 expression. CD3-APC/Cy7 (clone HIT3a), CD4-PE/Cy7 (clone a161A1), CXCR3-Pacific Blue (clone G025H7), CCR4-BV605 (clone L291H4), CCR6-FITC (clone G034E3), and CD71-APC (clone CY1G4). Compensation was conducted using single-stain PBMC controls. A third panel was used to measure activation and memory status after activation: CD62L-AlexaFluor488 (clone DREG-56), CD45RA-PECy5 (clone HI100) was used with the previous CD3, CD4, CD44 and CD71 antibodies.

For activation and culture of patients T cells, CD4 T cells were purified by negative selection using the human CD4 T Cell Isolation Kit (Miltenyi Biotec). T cells were activated with T cell activation/expansion CD2/3/28 beads at a ratio of 1:2 bead to cells (Miltenyi Biotec). Cultures were maintained in HPLM media with 5% dialyzed FBS (Sigma Aldrich) and supplemented with 100U/mL rhIL-2 from day 3 post-activation. For CD71 blockade experiments, cells were activated in the presence of either 2μg/mL anti-human CD71 (clone OKT9, BioXCell) or mouse IgG1 isotype control (clone MOPC-21).

### Cell cultures

Primary murine naïve CD4 T cells were isolated from the spleens and lymph nodes of SLE1.2.3 or control mice using the CD4+CD62L+ T cell isolation kit according to the manufacturer’s instructions (Miltenyi Biotec). CD62L positive selection was routinely ∼98% in purity as determined by flow cytometry. Naïve T cells were activated with irradiated splenocytes (irradiated at 30 Gy) in the presence of 1μg/ml anti-CD3. Unstimulated controls were incubated with mIL-7 at 0.1μg/ml. T cells were cultured at 37°C with 5% CO2 in RPMI 1640 media supplemented with 10mM HEPES, 50μM 2-mercaptoethanol, 100U/mL penicillin/streptomycin, and 10% heat inactivated FBS (Sigma Aldrich). Cultures were supplemented with recombinant human IL-2 (100U/mL) from day 3 post-activation. For differentiation of murine naïve CD4 T cells, media was supplemented with IL-12p70 (10ng/mL), IL-2 (100U/mL), anti-IL-4 (10μg/mL), anti-IFNγ (1μg/mL) for Th1 cells, or TGFβ (1.5ng/mL), IL-2 (100U/mL), anti-IL4 (10μg/mL), anti-IFNγ (10μg/mL) for iTreg cells. For human T cell differentiation, naïve CD4 T cells were isolated using the Naïve CD4 T Cell Isolation Kit II (Miltenyi). Human T cell Th17 cultures were activated with non-tissue culture treated plates with plate-bound anti-CD3 (3μg/mL), anti-CD28 (1μg/mL), and anti-ICOS (1μg/mL) antibodies. HPLM was supplemented with anti-IL-4 (2μg/mL), anti-IFNγ (2μg/mL), IL-23 (50ng/mL), IL-1β (50ng/mL), TGFβ1 (5ng/mL), IL-21 (25ng/mL) and IL-6 (40ng/mL). All cytokines were purchased from Peprotech except for IL-1β (R&D Systems).

The Plat-E retroviral packaging cell line (ATCC) was maintained at 37°C with 5% CO_2_ in DMEM media supplemented with 10% FBS, 100U/mL penicillin/streptomycin, 1μg/mL puromycin, and 10μg/mL blasticidin. Cell lines were tested for mycoplasma using the MycoProbe Mycoplasma Detection kit (R&D Systems).

### Flow cytometry

For intracellular and transcription factor stains, cells were first stained with viability dye +/- cell surface antibodies, fixed and permeabilized then stained for intracellular proteins with the appropriate kits. Transcription factor staining consisted of the eBioscience FoxP3/Transcription Factor Staining Buffer Set. For cytokine stains, cells were restimulated with 1μg/mL 12-myristate 13-acetate (PMA) and 750ng/mL ionomycin in the presence of GolgiPlug and GolgiStop for four hours, then processed as other intracellular stains. Unstimulated cells served as a negative control. Antibodies for IL-10-producing T cells: CD4-e450 (clone GK1.5) and IL-10-PE (clone JES5-16E3). B cells were quantified with B220-e450 (clone RA3-6B2) and CD19-APC (clone 6D5). Treg and Tfr cells were quantified with CD4-PECy7, CD25-APC (clone PC61.5), FoxP3-e450 (clone FJK-16s) and Bcl-6-FITC (clone 7D1). Activation markers used: CD25-PE, CD44-PECy5 (clone IM7), CD69-FITC (clone H1.2F3), CD62L-APC (clone MEL-14), and phospho-S6 (Ser235,236)-APC (clone cupk43k).

For mitochondrial dyes, Mitotracker Green and MitoSOX Red were combined with cell surface CD4-e450 and CD71-APC antibody in complete media at 37°C with 5% CO_2_ for 25 minutes. Labile iron staining was conducted with BioTracker Far Red Labile 2+ Live Cell Dye (Millipore). Cells were first washed with Hank’s balanced salt solution (HBSS), and then stained for 30 minutes at 37°C with 5% CO_2_ in a 5μM solution of dye in HBSS. Unstimulated T cells and T cells with no dye were used as negative controls.

### CRISPR/Cas9 Screens

Details for library design and preparation were previously described^60^. Briefly, guide RNA (gRNA) sequences were chosen from the Brie library. Four gRNA sequences for each gene and five non-targeting controls (NTCs), flanked by adapter sequences, were purchased as an oligo pool from Twist Bioscience. The iron metabolism library contained 55 total gene targets and 225 total gRNAs. The pMx-U6-gRNA-GFP vector was previously modified to express BFP in the place of GFP, and oligos were cloned into this vector using Gibson Assembly Master Mix. DNA was packaged into retrovirus following transfection of the Plat-E retroviral packaging cell line. Retroviral transduction of primary T cells was performed with retronectin-coated plates (Takara Bioscience). Naïve T cells had been activated as described in ‘Cell Cultures’ methods for two days before transduction. CRISPR/Cas9 screens were conducted in HPLM media^61^ and supplemented with 100U/mL rhIL-2 after transduction.

Collection and analysis of the gRNA frequencies over time followed the same methodology as previously described by Sugiura *et al*.^60^. At least 1,000-fold representation of the library was maintained throughout the process of cell harvesting, amplification and sequencing. FASTQ files were analyzed using the Model-based Analysis of Genome-wide CRISPR/Cas9 Knockout (MAGeCK v0.5.0.3) method to determine statistically significant gRNA enrichments or depletions.

### ELISAs

Autoantibodies were measured with ELISA kits purchased from Alpha Diagnostics: Anti-Histone Total Ig, Anti-Nuclear Antibodies Total Ig, and Anti-dsDNA Total Ig. Serum was diluted 1:500 and samples run in technical duplicates. Optical density at 450nm was measured on the Cytation5 Imager. For cytokine measurements, serum was subjected to the LEGENDplex Mouse Inflammation Panel assay (BioLegend). Technical duplicates were averaged and compared to a standard curve according to the manufacturer’s protocol. IL-10 in human cell culture supernatants were measured using the LEGEND Max Human IL-10 ELISA kit (BioLegend). Technical duplicates were averaged and then normalized to the total number of live cells in each sample at the time of collection.

### Extracellular Flux Analyses

Extracellular flux analysis was performed with the Seahorse XFe96 Analyzer (Agilent). Plates were coated with Cell-Tak solution (Corning) for 30 minutes at room temperature before seeding the cells. 150,000 viable cells were seeded per well and a minimum of five technical replicates were seeded for each sample. Final cell counts of each well were acquired by bright field imaging using a Cytation5 imager for normalization. The Glycolysis Stress Test was performed according to the Agilent protocol recommendations. The Mito Stress Test was performed with oligomycin A at 1.5μM, FCCP at 1.5μM, and rotenone/antimycin A at 0.5μM final concentrations.

### ICP-MS

To measure intracellular iron concentrations, equal cell numbers were determined and placed into metal-free 15 mL conicals (VWR). Cell pellets were washed twice with 10 mL of PBS and then stored at -80°C until downstream processing. Pellets were digested in 200 μL Optima-grade nitric acid (Fisher) and 50 μL Ultratrace-grade hydrogen peroxide (Sigma) and incubated overnight at 65°C. The next day, 2 mL of ultrapure-grade water (Invitrogen) was added before analysis.

Elemental quantification on acid-digested samples was performed using an Agilent 7700 inductively coupled plasma mass spectrometer (Agilent, Santa Clara, CA) attached to a Teledyne CETAC Technologies ASX-560 autosampler (Teledyne CETAC Technologies, Omaha, NE). The following settings were fixed for the analysis Cell Entrance = -40 V, Cell Exit = -60 V, Plate Bias = -60 V, OctP Bias = -18 V, and collision cell Helium Flow = 4.5 mL/min. Optimal voltages for Extract 2, Omega Bias, Omega Lens, OctP RF, and Deflect were determined empirically before each sample set was analyzed. Element calibration curves were generated using ARISTAR ICP Standard Mix (VWR, Radnor, PA). Samples were introduced by peristaltic pump with 0.5 mm internal diameter tubing through a MicroMist borosilicate glass nebulizer (Agilent). Samples were initially up taken at 0.5 rps for 30 s followed by 30 s at 0.1 rps to stabilize the signal. Samples were analyzed in Spectrum mode at 0.1 rps collecting three points across each peak and performing three replicates of 100 sweeps for each element analyzed. Sampling probe and tubing were rinsed for 20 s at 0.5 rps with 2% nitric acid between every sample. Data were acquired and analyzed using the Agilent Mass Hunter Workstation Software version A.01.02.

### Histology and Immunohistochemistry

HE sections were scored in a semi-quantitative fashion by a pathologist^41^. For liver sections, perivascular inflammatory cell infiltrates were scored on a scale from 0-3 where 0 = no pathology, 1 = mild, rare, few inflammatory cells, 2 = multifocal to coalescing zones of inflammatory cells and 3 = severe, coalescing to diffuse inflammatory infiltrate. In the kidney sections interstitial inflammation was scored using the same scoring system as described for the liver. Additional sections were cut for immunohistochemistry for both CD4 and F4/80 which were stained using the Leica Bond Rx automated immunostainer (Leica, Buffalo Grove, IL). Briefly, slides were deparaffinized for both assays and antigen retrieval was performed using Proteinase K (Dako, Agilent, Santa Clara, CA) for F4/80 for 7 minutes at room temperature and EDTA at pH of 9.0 (Leica, Buffalo Grove, IL) for CD4 for 20 minutes on the immunostainer. Slides were blocked with Leica Block (Leica) for 10 minutes at 37°C for F4/80 or blocked with a protein block (catalog number x0909, Dako) for 10 minutes followed by an Fc block (catalog number 553142, BD Pharmingen, Franklin Lakes, NJ) at 1:1,000 for 15 minutes at 37°C. Slides were incubated with anti-F4/80 (NB600-404, Novus Biologicals LLC, Littleton, CO) for one hour at a 1:600 dilution and then incubated in a rabbit anti-rat secondary (BA-4001, Vector Laboratories, Inc., Burlingame, CA) for 12mins at a 1:7500 dilution. Slides were incubated with anti-CD4 (ref# HS-360 017, SYSY, Goettingen, Germany) for 60minutes at a 1:1500 dilution and followed by a biotinylated anti-rat (Cat. BA-5000, Vector Laboratories, Inc.) for 15 minutes at a 1:2000 dilution. The Bond Polymer Refine detection system was used for visualization for both assays. Slides were then dehydrated, cleared and coverslipped.

### Transmission electron microscopy

T cells were activated as described previously and supplemented with IL-2 on day 3. On day 5, cells were washed with warm PBS and then fixed in 2.5% glutaraldehyde in 0.1 M cacodylate for 1 hour at room temperature followed by 24 hours at 4°C. Following fixation, the cells were postfixed in 1% OsO4 and en bloc stained with 1% uranyl acetate the dehydrated in a graded ethanol series. Samples were gradually infiltrated with Quetol 651 based Spurr’s resin with propylene oxide as the transition solvent. The Spurr’s resin was polymerized at 60°C for 48 hours. Blocks were sectioned on a Leica UC7 ultramicrotome at 70 nm nominal thickness and the samples stained with 2% uranyl acetate and lead citrate. Transmission electron microscopy was performed using a Tecnai T12 operating at 100 kV with an AMT NanoSprint CMOS camera using AMT imaging software for single images and SerialEM for tiled datasets. Tiled datasets were reconstructed using the IMOD/eTomo software suite. Mitochondria quantification was done in FIJI by manually segmenting all mitochondria within cell cross sections from the tiled TEM datasets until at least 100 mitochondria were measured. Mitochondria area fraction was determined by summing the cross-sectional area of all mitochondria within a cell and divided by the area of the entire cell.

### Statistics

Statistical analyses were performed with GraphPad Prism software (v9). Statistically significant results are labelled as follows: * p < 0.05, ** p < 0.01, *** p < 0.001, **** p ≤ 0.001. For a comparison of two groups, student’s *t*-test was performed. For more than two groups, one-way ANOVA was performed. Figures with data points connected by lines are indicative of paired analyses, whereas all other statistical tests were unpaired. In all figures, error bars represent the mean ± standard deviation.

## Acknowledgements

We thank Annette Oeser for collecting patient samples. Brenna D. Appleton and Megan Dickson for colony maintenance of lupus mice. Katherine L. Beier and Matthew Z. Madden for technical assistance on certain experiments. K.V. would like to thank Jackie E. Bader for helping with *in vivo* CD71 experiments as needed. We acknowledge the Translational Pathology Shared Resource supported by NCI/NIH Cancer Center Support Grant 5P30 CA68485-19 and the Shared Instrumentation Grant S10 OD023475-01A1 for the Leica Bond RX used for immunohistochemistry.

TEM was performed in part through the use of the Vanderbilt Cell Imaging Shared Resource (supported by NIH grants CA68485, DK20593, DK58404, DK59637 and EY08126). This work was supported by the William Paul Distinguished Innovator for the Lupus Research Alliance (J.C.R.), R01DK105550 (J.C.R.), T32 DK101003 (K.V.), R01AI150701 (E.P.S.).

## Author Contributions

K.V. conceptualized the study, designed all experiments, conducted experiments and analyses, and wrote the first draft of the manuscript. A.C.Y, A.E.S., and J.H.B performed some experiments and provided technical assistance. K.N.GC. analyzed, scored and interpreted histology of lupus mice. E.S.K. performed, analyzed and interpreted electron microscopy experiments. A.S. designed the CRISPR system used for screening. W.N.B. performed ICP-MS experiments and was supervised by E.P.S. M.J.O. provided the IRB to acquire lupus patient samples, calculated SLEDAI scores, and supervised clinical aspects of the manuscript as needed. A.S.M. provided lupus mice, overall assistance, and intellectual contributions. J.C.R. supervised the project and research design and writing the manuscript.

---

**Supplemental Fig. 1:**
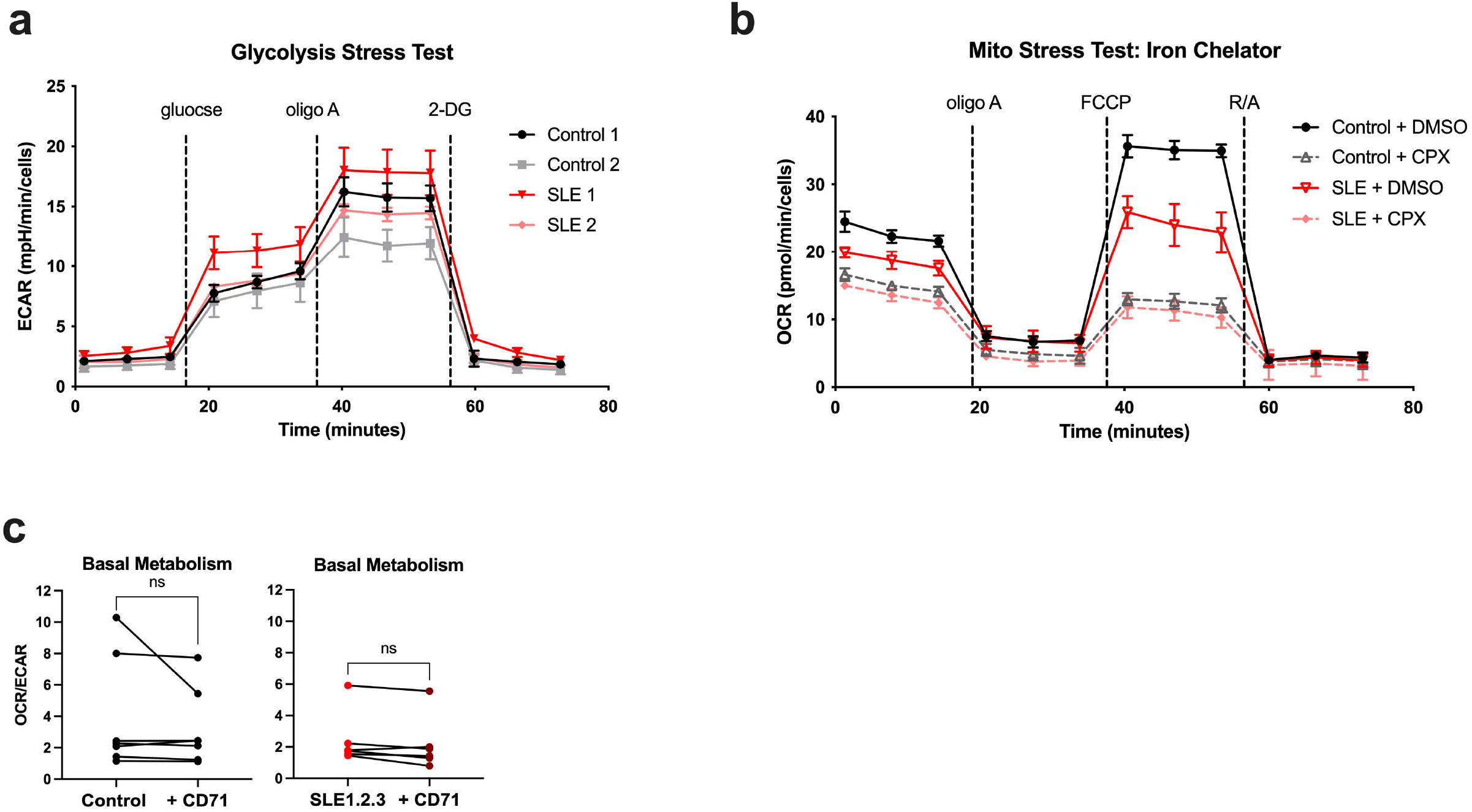
Extracellular flux analyses of SLE1.2.3 T cells. **a,** Glycolysis Stress Test is shown from two biological replicates. **b,** On day 5 post-activation, T cell cultures were incubated with the iron chelator ciclopirox olamine (CPX) for 4 hours and then subjected to Mito Stress Test. Representative of two independent experiments **c,** Basal metabolism was determined from basal OCR and ECAR in the Mito Stress Test as described in Fig. 4d. Paired *t*-test, ns= not significant.

**Supplemental Fig. 2:**
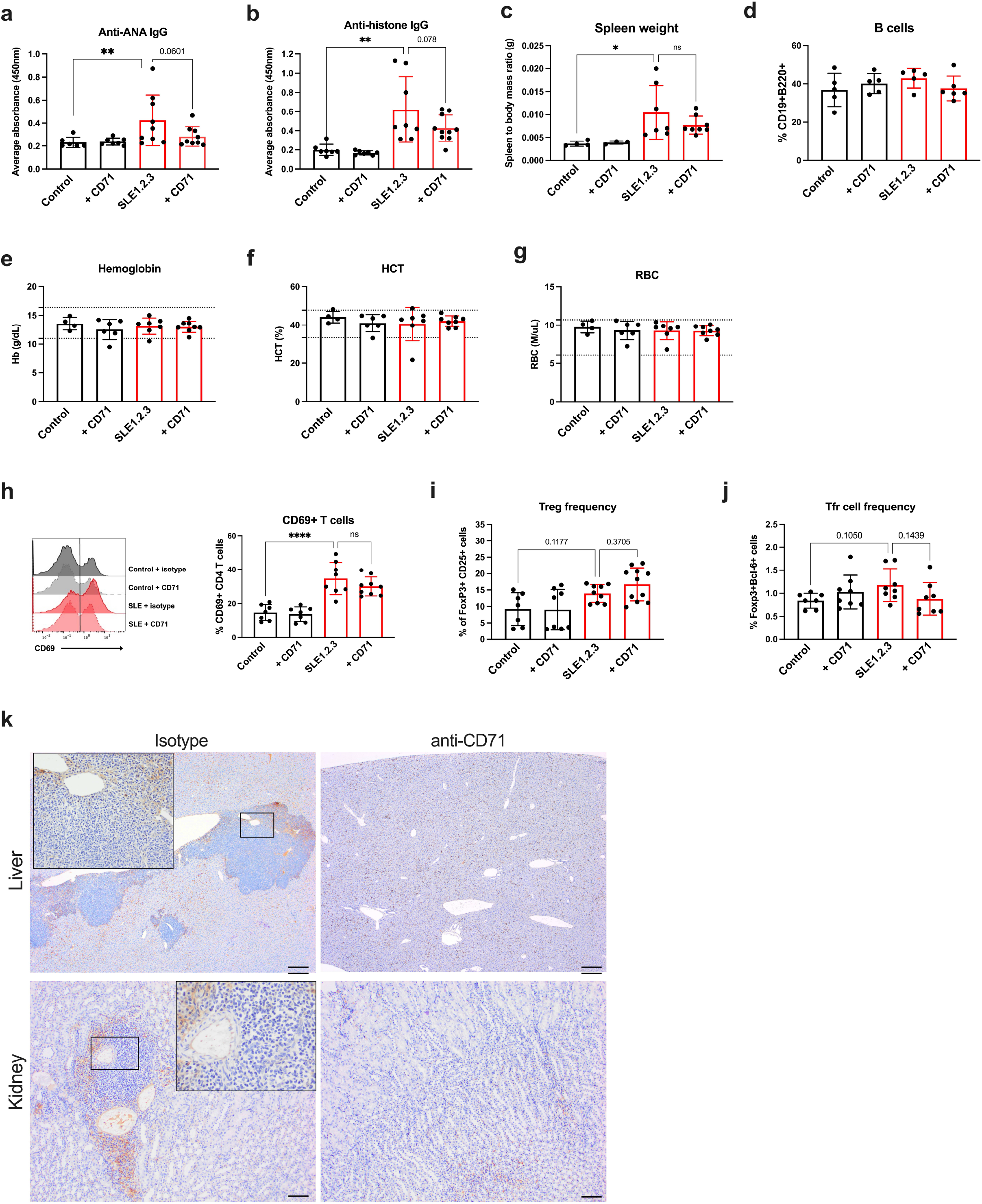
*In vivo* CD71-blockade in SLE1.2.3 mice. **a,b,** Endpoint serum was analyzed for anti-nuclear antibodies (ANA), and anti-histone IgG antibodies (b). **c,** Spleen weight divided by body mass. **d,** Percentages of B cells within splenocyte population were quantified by double positive CD19 and B220. (N=2 experiments) **e,** Complete blood count (CBC) results for hemoglobin, **f,** hematocrit (HCT), and **g,** red blood cell (RBC) counts. Normal ranges from B6 mice are indicated by the dotted lines. **h,** Endpoint CD4 T cells were subjected to flow cytometry to determine CD69 expression. **i,j,** Spleens and lymph nodes were examined for Treg (i) and T follicular regulatory (Tfr) (j) cell frequencies at the study endpoint.. **k,** F4/80 staining representative IHC images from the liver and kidney. Scale bar= 200μm. Similar results were obtained in three mice per treatment group. (a,b,c,h,i,j) One-way ANOVA with Sidak’s multiple comparisons test.

**Supplemental Fig. 3:**
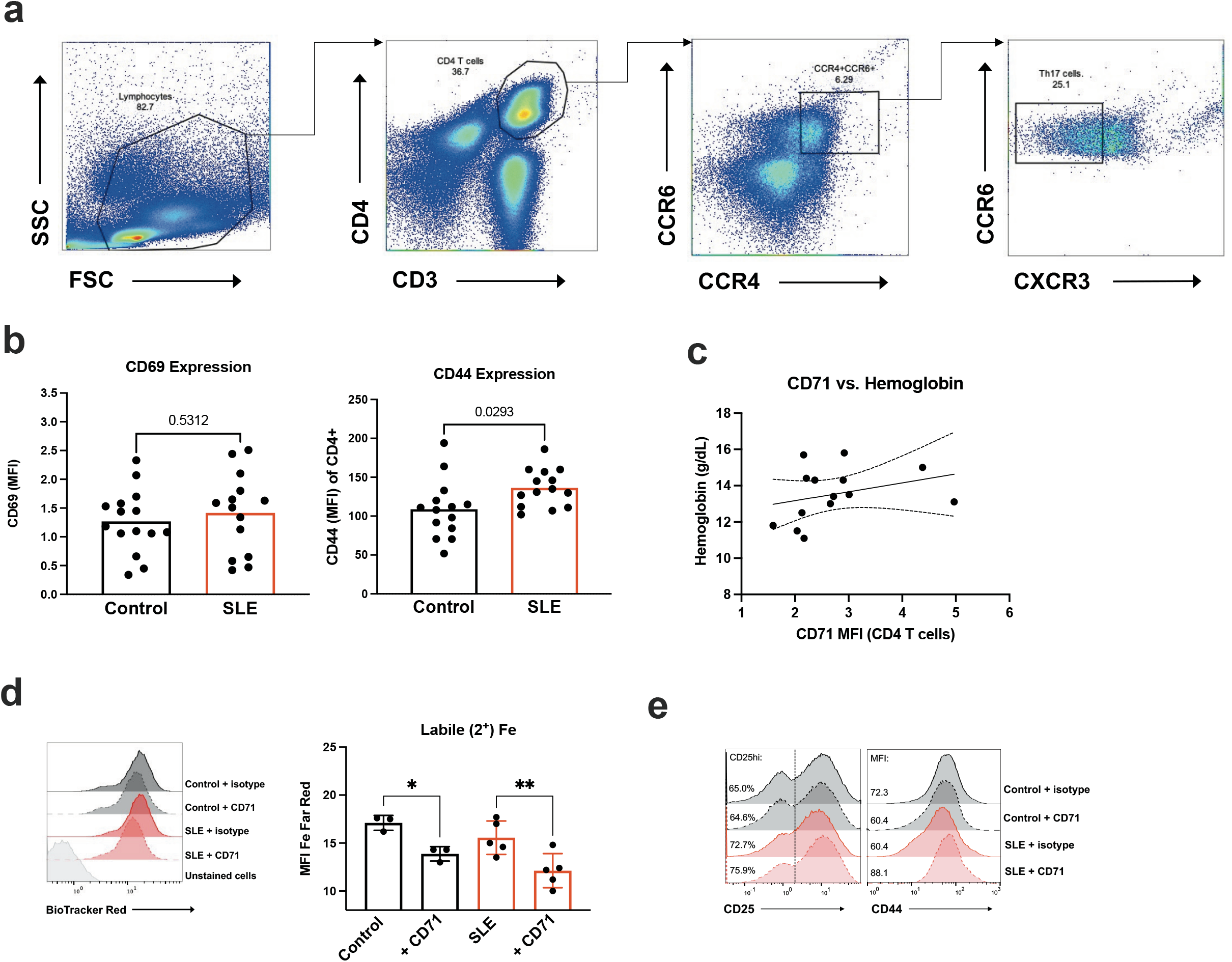
Supporting SLE patient data. **a,** Gating strategy for PBMC flow cytometry experiments, showing identification of Th17 subset. **b,** CD69 and CD44 expression on bulk CD4 T cells from PBMC samples. Student’s t-test. **c,** SLE patient CD71 expression on CD4 T cells compared to hemoglobin levels. **d,** Biotracker Red labile iron dye staining on day 3 post-activation. One-way ANOVA with Sidak’s multiple comparisons test. **e,** Representative activation marker staining on day 3 post-activation +/- CD71 blockade. Similar results were obtained in at least 8 independent experiments.

**Supplemental Table 1:** SLE Patient information.

|  | SLE<br>Median [IQR], or # (%) |
| --- | --- |
| Age, years | 34 [29, 40] |
| Sex, # female | 12 (75) |
| Race or ethnicity |  |
| White, # (%) | 9 (56) |
| Black, # (%) | 2 (13) |
| Asian, # (%) | 1 (6) |
| Hispanic/Latino, # (%) | 2 (13) |
| ANA positivity, # (%) | 16 (100) |
| Anti-dsDNA positivity, # (%) | 12 (75) |
| Renal involvement, ever # (%) | 8 (50) |
| SLEDAI, score | 4 [2, 5] |
| SLICC, score | 1 [0, 1] |
| Current hydroxychloroquine use, # (%) | 16 (100) |
| Current corticosteroid use, # (%) | 4 (33) |
| Current mycophenolate mofetil use, # (%) | 9 (56) |

IQR= interquartile range. ANA= antinuclear antibody. Anti-dsDNA= anti-double stranded DNA antibody. SLEDAI = SLE disease activity index. SLICC = Systemic Lupus International Collaborating Clinics damage index.

